# Nitrogen cycling during an Arctic bloom: from chemolithotrophy to nitrogen assimilation

**DOI:** 10.1101/2024.02.27.582273

**Authors:** Rafael Laso Pérez, Juan Rivas Santisteban, Nuria Fernandez-Gonzalez, Christopher J. Mundy, Javier Tamames, Carlos Pedrós-Alió

## Abstract

In the Arctic, phytoplankton blooms are recurring phenomena occurring during the spring-summer seasons and influenced by the strong polar seasonality. Besides, bloom dynamics is affected by nutrient availability, especially nitrogen, which is the main limiting nutrient in the Arctic. This study aimed to investigate the changes in an Arctic microbial community during a phytoplankton bloom with a special focus on the nitrogen cycle. Using metagenomic and metatranscriptomic samples from the Dease Strait (Canada) from March to July (2014), we reconstructed 176 metagenome-assembled genomes. Bacteria dominated the microbial community, although archaea reached up to 25% of genomic abundance in early spring, when *Nitrososphaeria* archaea actively expressed genes associated with ammonia oxidation to nitrite (*amt, amoA, nirK*). The resulting nitrite was presumably further oxidized to nitrate by a *Nitrospinota* bacterium that highly expressed a nitrite oxidoreductase gene (*nxr*). Since May, the constant increase in chlorophyll *a* indicated the occurrence of a phytoplankton bloom, promoting the successive proliferation of different groups of chemoorganotrophic bacteria (*Bacteroidetes*, *Alphaproteobacteria* and *Gammaproteobacteria*). These bacterial taxa showed different strategies to obtain nitrogen, whether it be from organic or inorganic sources, according to the expression patterns of genes encoding transporters for nitrogen compounds. In contrast, during summer, the chemolithotrophic organisms thriving during winter, reduced their relative abundance and the expression of their catabolic genes. Based on the functional analysis of our data, we see a transition from a community where nitrogen-based chemolitotrophy plays a relevant role, to a chemoorganotrophic community based on the carbohydrates released during the phytoplankton bloom, where different groups specialize in different nitrogen sources.

## Introduction

The Arctic region is a unique environment characterized by extreme seasonal transitions between ice-covered winters and ice-free summers. The stress of these transitions has increased due to climate change affecting the whole ecosystem: from physicochemical variables to biological interactions. Scientists have observed an increase of phytoplankton biomass with subsequent higher net primary production (NPP) owing to the sea ice decline and a larger supply of nutrients into the Arctic system^1^. Primary production is associated to recurrent phytoplankton blooms during the summer months, which are dependent on the availability of nutrients like nitrogen or phosphorus. Nitrogen is the primary nutrient affecting phytoplankton growth in the Arctic, since it becomes limiting after the spring-summer bloom^2–5^. In recent years, the biogeochemistry of the Arctic nitrogen cycle and its relationship to primary productivity has been studied in more detail, showing that nitrogen derived from rivers and coastal erosion supports around 28-51% of the Arctic NPP^6^ or that nitrogen fixation might be a more important process than previously thought in ice-free waters^7^. By using 16S rRNA gene surveys, previous studies linked different taxonomic groups with specific nitrogen compounds^8^. Now, the emergence of metagenomics has opened a new perspective to investigate the function of prokaryotes in the cycling of nitrogen. Some studies focused on the role of certain groups on specific processes like the diversity of Arctic diazotrophs^9^ or the use of urea in nitrification by polar *Thaumarchaeota*^10^. Fewer studies have investigated nitrogen cycling at a community perspective. One example is the work of Royo-Llonch et al. (2021), who showed that chemolithoautotrophic metabolisms like ammonia or nitrite oxidation are more prevalent during spring and autumn, but in general they are not so widespread as heterotrophic or mixotrophic lifestyles^11^. In a different study, the presence of nitrate-reducing genes in different polar MAGs was understood as a “strategy of metabolic versatility to adapt to environmental change” ^12^. In the Canadian Arctic Archipelago, a metagenomics study captured spatial differences between seawater and sea ice in relation to different functions, like nitrate and nitrite, although these differences were not significant^12^. Here, we want to better characterize the prokaryotic community during the seasonal winter-summer transition. Concretely, we have investigated the role of the prokaryotic community in the cycling of nitrogen by linking genomic information with expression data and studying how the functional profile changed from a winter scenario to a summer bloom situation. For this purpose, we collected water samples from the Dease Strait from March to July in 2014.

## Results

### Community composition

Using 13 metagenomic samples retrieved from the Dease Strait (Canadian Arctic, Figure 1a) from March until July 2014 (Table S1), we reconstructed 176 metagenome-assembled genomes (MAGs). We also collected samples for metatranscriptomic analysis. During the sampling period, a phytoplankton bloom developed as shown by the increase in chlorophyll *a* concentration (Figure 1b). Coverage analysis showed that our MAGs captured approximately half of the metagenomic reads across all samples (Table S2). According to the genomic relative abundance (Figure 1c; Table S2), the dominant groups were similar to those detected in previous 16S rRNA gene surveys in the Arctic Archipelago: ammonia-oxidizing *Thaumarchaeota* for archaea^13,14^ and the bacterial groups *Alphaproteobacteria, Gammaproteobacteria* and *Flavobacteriaceae* (*Bacteroidetes*)^15–19^. Bacteria dominated during the whole period, although archaea reached up to 25% of the prokaryotic community in March, when a *Nitrososphaerales* MAG was the most abundant organism of the whole dataset (19-21%) and decreased in abundance during the spring, a pattern that has been previously reported for polar *Nitrososphaerales*^10^. In contrast, archaeal MAGs from the order *Poseidonales* showed lower relative abundances, although some seasonal changes were observed with higher relative abundances during winter-spring (4-6% vs. 2-3% in summer). For bacteria, several groups had a strong seasonality presumably linked to the phytoplankton bloom. For instance, *Flavobacteriales* MAGs (class *Bacteroidia*) represented 7% of the prokaryotic assemblage in March and reached 57% in late July, at the peak of the bloom. This seasonality is linked to the metabolic lifestyle of certain marine *Flavobacteriales*, which are specialized in degrading phytoplankton-derived carbohydates^20^. Other groups showing seasonal changes were the alphaproteobacterial *Pelagibacterales* and the gammaproteobacterial *Pseudomonadales*, PS1 (that includes the family *Thioglobaceae*) and SAR86. The gammaproteobacterial groups increased their abundance during May to peak in June and decreased during the last period of the sampling. In contrast, the alphaproteobacterial *Pelagibacterales* slowly decreased in their relative abundance during middle spring to increase in abundance dramatically at the end of July (27.5%). These changes were mostly associated with a single MAG (Alphaproteobacteria_10), which always showed a high relative abundance (always >6%).

**Figure 1.**
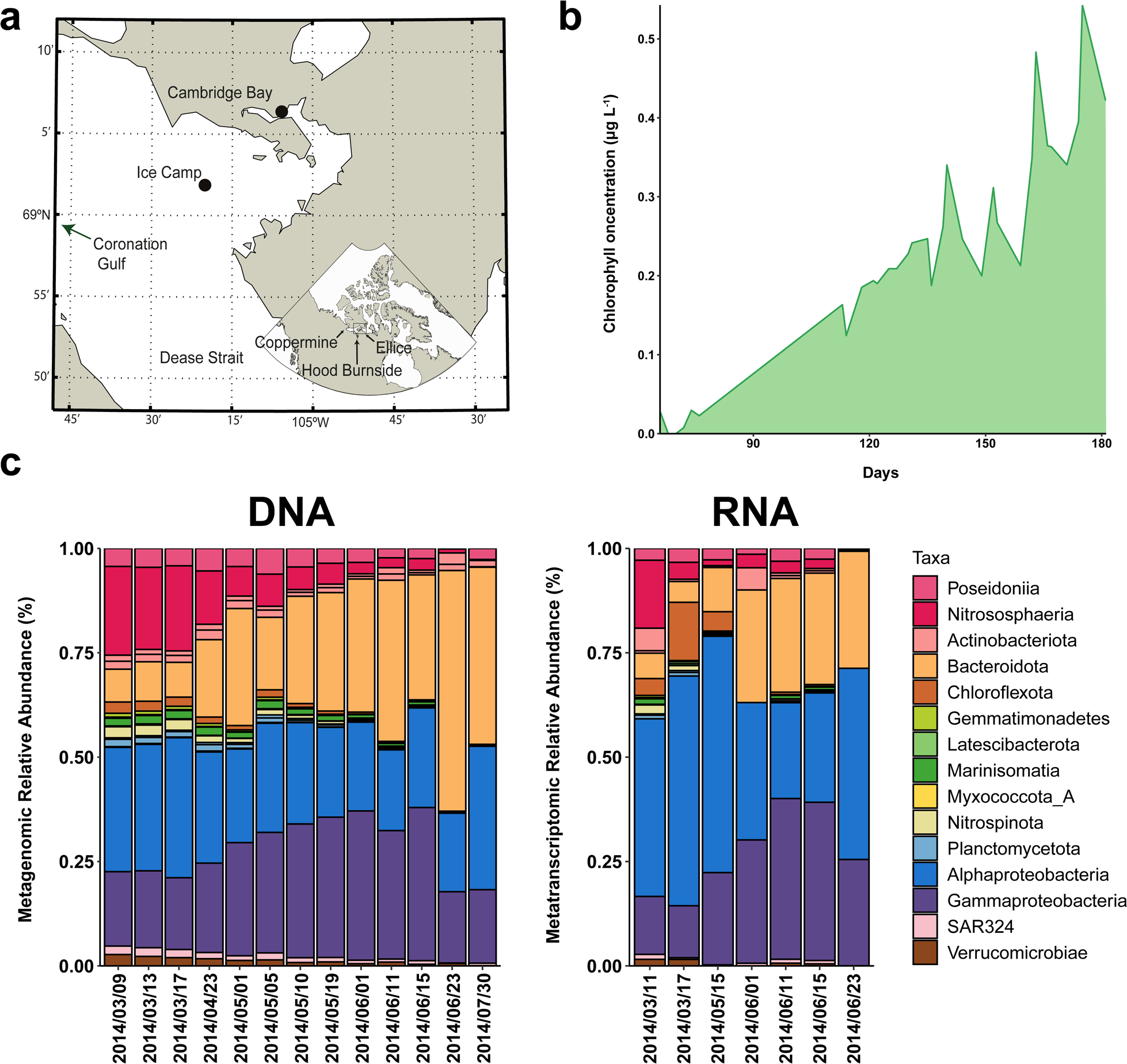
Overview of the sampling site, the bloom conditions and the MAG abundance. **A** Map of the sampling site. **B** Graph showing the chlorophyll concentration over time. The increase in chlorophyll indicates a phytoplankton bloom. **C** Relative abundance at the DNA and RNA level of our MAG dataset grouped by class.

Metatranscriptomic analysis showed similar trends at higher taxonomic levels: higher expression levels of archaea in winter-spring, increased expression levels for *Flavobacteriales* and *Gammaproteobacteria* at late spring, and a high relative expression across the whole period for *Alphaproteobacteria*. Most of the transcriptomic reads mapped to a few MAGs in all the samples. For instance, the abundant *Pelagibacterales* MAG (Alphaproteobacteria_10) represented always more than half of the RNA reads mapping to the whole *Alphaproteobacteria*. In fact, this MAG captured more than half of the mapped reads during mid-March and May and its lowest transcriptomic relative abundance was 12% in mid-June. During winter, the only *Nitrosopumilus* MAG (Nitrosopumilus_01) reached 16% of transcriptomic relative abundance. In contrast, two gammaproteobacterial MAGs, a PS1 (Gammaproteobacteria_11) and a *Pseudomonadales* (Gammaproteobacteria_22) increased their transcriptomic relative abundance from 5% and <1% respectively in winter to >10% in May-June. In *Bacteroidetes,* no MAG reached relative abundances over 10%, but still two *Flavobacteriales* peaked at 5-6% abundance in May-June. Finally, it is notable that two groups, *Actinomycetia* and *Chloroflexota* reached considerable transcriptomic relative abundances (5% and 13% respectively) during winter-spring in contrast to their metagenomic relative abundances, which was always below 3%.

### Functional analysis

To study the nitrogen cycling within our dataset, we searched in each MAG for key genes of the different nitrogen cycling processes using specific profiles (see Material and Methods). The retrieved genes were then classified with a phylogenetic database that allows to distinguish the nitrogen cycling genes from closely related paralogs. The taxonomic composition and expression levels of the nitrogen-cycling genes showed a succession from a community based on ammonia chemolithotrophy to another one based on chemoorganotrophy due to the algal bloom (Figure 2, Table S3).

**Figure 2.**
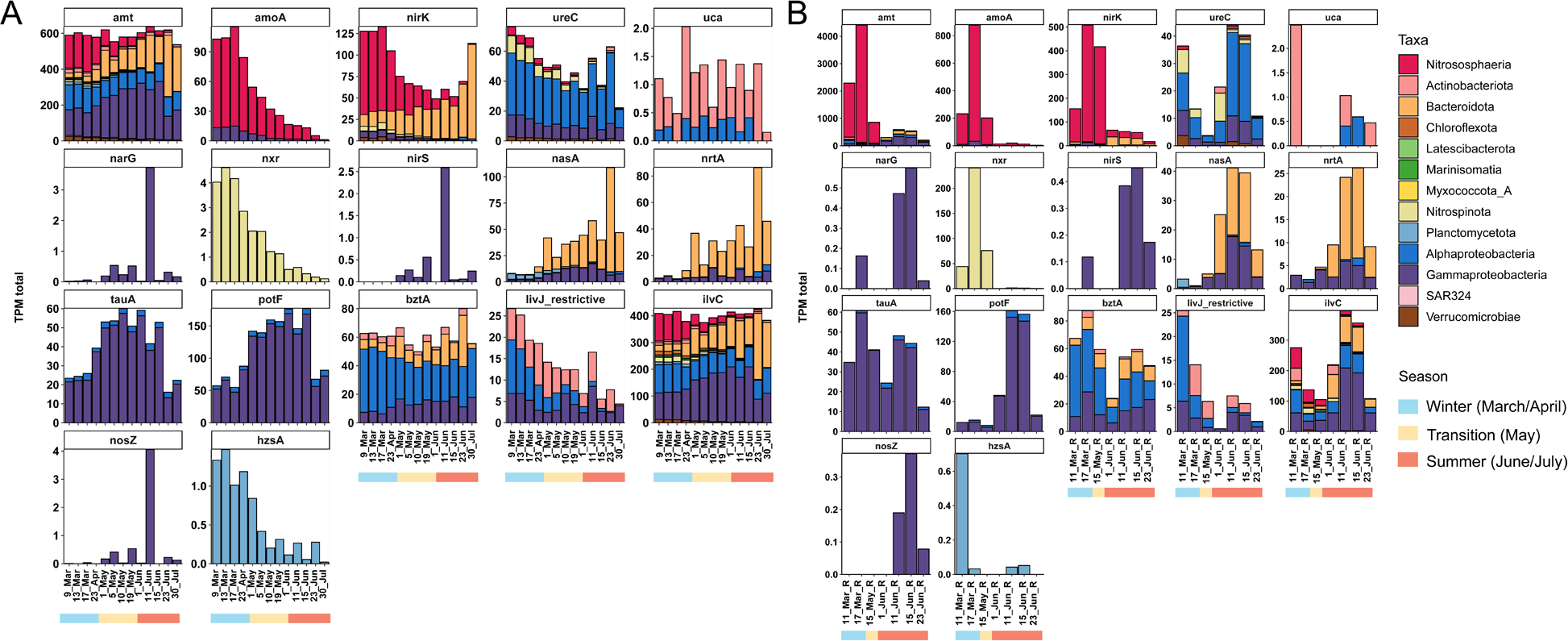
General overview of the nitrogen-cycling genes of our MAG dataset and their expression. **A** Relative DNA abundances profiles, based on TPM, of the different genes grouped by taxonomical class. **B** Total RNA TPM profiles of the different genes grouped by taxonomical class.

### Ammonia oxidation by archaea is prevalent during winter-early spring

In the marine environment, the process of ammonia oxidation is mostly performed by ammonia-oxidizing archaea (AOA) from the class *Nitrososphaeria* (previously known as *Thaumarchaeota*), although there are bacterial groups with the same metabolism. The key enzyme of this process is the ammonia monooxygenase (Amo). Two MAGs in our dataset, a *Nitrososphaeria* (Nitrosopumilus_01) and a *Burkholderiales* (Gammaproteobacteria_04), encoded an Amo enzyme, but only the *Nitrososphaeria* showed high genomic and transcriptomic relative abundances in the dataset. Specifically, during winter the *Nitrososphaeria* had extremely high transcriptomic levels, with an expression peak for the *amoA* gene (which encodes the main catalytic subunit from Amo) in mid-March, followed by a decrease and almost no expression signal in June (Figure S1). Other *Nitrososphaeria* genes involved in ammonia oxidation showed a similar transcription pattern like the *amt* gene, which codes for the transmembrane ammonium transporter, or the gene encoding NirK, a nitrite reductase (Figure S1). The NirK enzyme usually catalyzes the reduction of nitrite (NO ^-^) to nitric oxide (NO), but in archaea it was proposed that NirK could substitute the function of the canonical hydroxylamine oxidoreductase (hao), which in ammonia-oxidizing bacteria is responsible for the second step of the ammonia oxidation, but it is absent in all ammonia-oxidizing archaea^10,11^. The similar transcriptomic profile for the *amoA*, *amt* and *nirK* genes of the *Nitrososphaeria* MAG indicates that this organism thrives during winter by oxidizing ammonia. Similar results have been observed for Arctic AOA^12–15^, which perform ammonia oxidation during winter due to the high ammonia concentrations and disappear during summer because of the ammonia depletion and a potential photoinhibition^16^.

### A *Nitrospinota* bacterium oxidizes nitrite during the winter and early-spring

We recovered three MAGs affiliated to the bacterial class *Nitrospinia* within the phylum *Nitrospinota.* Although the three MAGs together did not reach above 3% relative abundance, one of them affiliated to the LS-NOB genus (Nitrospinia_01) has the genes encoding for a nitrite oxido-reductase (Nxr) enzyme, which show intense transcription during winter (Figure S1). This enzyme, responsible for the oxidation of nitrite to nitrate, is a hallmark for most *Nitrospinia*^17^, although closely related homologues (nitrate reductase, Nar) are present in other organisms reducing nitrate to nitrite^18^. Our phylogenetic analysis resolved that the *Nitrospinota* LS-NOB genome encodes a true Nxr, which has an expression peak during winter, similar to the genes of the ammonia oxidizing archaea. This suggests a metabolic link between the ammonia-oxidizing *Nitrososphaeria* and the nitrite-oxidizing *Nitrospinota*. Specifically, the oxidation of ammonia by the archaeon produces nitrite as metabolic end product, which can then be used for nitrite oxidation by the *Nitrospinota* MAG. Therefore, the expression patterns of both organisms are really similar and when summer arrives and the *Nitrososphaeria* MAG decreases in abundance, the expression of the *Nitrospinota nxr* also drops, probably due to the decrease on nitrite (Figure S1). This co-occurrence of *Nitrososphaeria/Nitrospinota* has been previously observed in Arctic waters^12,19^. There were two additional *Nitrospinia* MAGs affiliated to the genus *Nitromaritima* (Nitrospinia_02 and Nitrospinia_03), in which we could not detect genes for the Nxr complex. The absence of these genes is surprising, since the Nxr complex is a hallmark of most *Nitrospinia*^17^ and this complex is encoded in other *Nitromaritima* genomes^20^. This absence might be attributed to the binning process since both genomes have completeness levels below 60%. Still, both genomes encoded for a urease enzyme, absent in the LS-NOB *Nitrospinota* MAG. The *ureC* gene of Nitrospinia_03 showed moderate expression levels in March and early June and could indicate that these *Nitrosomaritima* are using urea as nitrogen and energy source, instead of oxidizing nitrite (Table S3). Genes encoding ureases have been detected in several *Nitrospinia* genomes^20,21^, where they could serve to degrade urea to ammonia, which could then be used by ammonia-oxidizing organisms.

### Heterotrophic bacteria thriving in summer can use different nitrogen sources

With the winter-summer transition, there is a shift in the community with the growth and decay of specific groups due to the seasonal changes (i.e. melting of sea-ice, increase in solar irradiance) and the phytoplankton bloom^22^. The majority of the spring-summer microorganisms are heterotrophic bacteria feeding on the carbohydrates released by the phytoplankton^22^. These organisms still require a nitrogen source for their growth, developing a competition for the different nitrogen compounds reflected in the metatranscriptomic profiles (Figure 3). Most MAGs of our dataset have genes for an ammonium transporter (Amt, Figure 4), which are highly expressed during the summer phytoplankton bloom (Figure 3). During winter the archaeal *amt* genes represented 90% of all *amt* transcripts (Figure 2b), but at the beginning of June there is a shift in the transcriptomic profile and the archaeal *amt* expression plummets, while the bacterial *amt* genes are highly transcribed, especially those of the most abundant groups: *Gammaproteobacteria, Alphaproteobacteria* and *Bacteroidota.* Although the widespread and high transcription of *amt* genes indicates a preference for ammonia as nitrogen source, many of the MAGs of our dataset possess genes to incorporate nitrogen from different sources, indicating a diversity of strategies probably due to strong competition for nitrogen during the phytoplankton bloom. Interestingly, we could not find any organisms with the ability to fix nitrogen gas. No nitrogenase genes were found in the whole dataset. Although diazotrophy is becoming a more prominent metabolism in the Arctic environment^31,32^, metagenomics studies are unable to detect the presence of the nitrogen-fixing genes^33^ or even *nifH* gene surveys do not always detect them in different kind of environments^34^. Therefore, the absence of *nifH* genes in our dataset could be due to the absence of diazotrophic organisms or because they are really low in abundance. Additionally, we could detect other genes to incorporate alternative nitrogen sources including inorganic (nitrate) and organic (taurine, putrescine, amino acids, urea) compounds.

**Figure 3.**
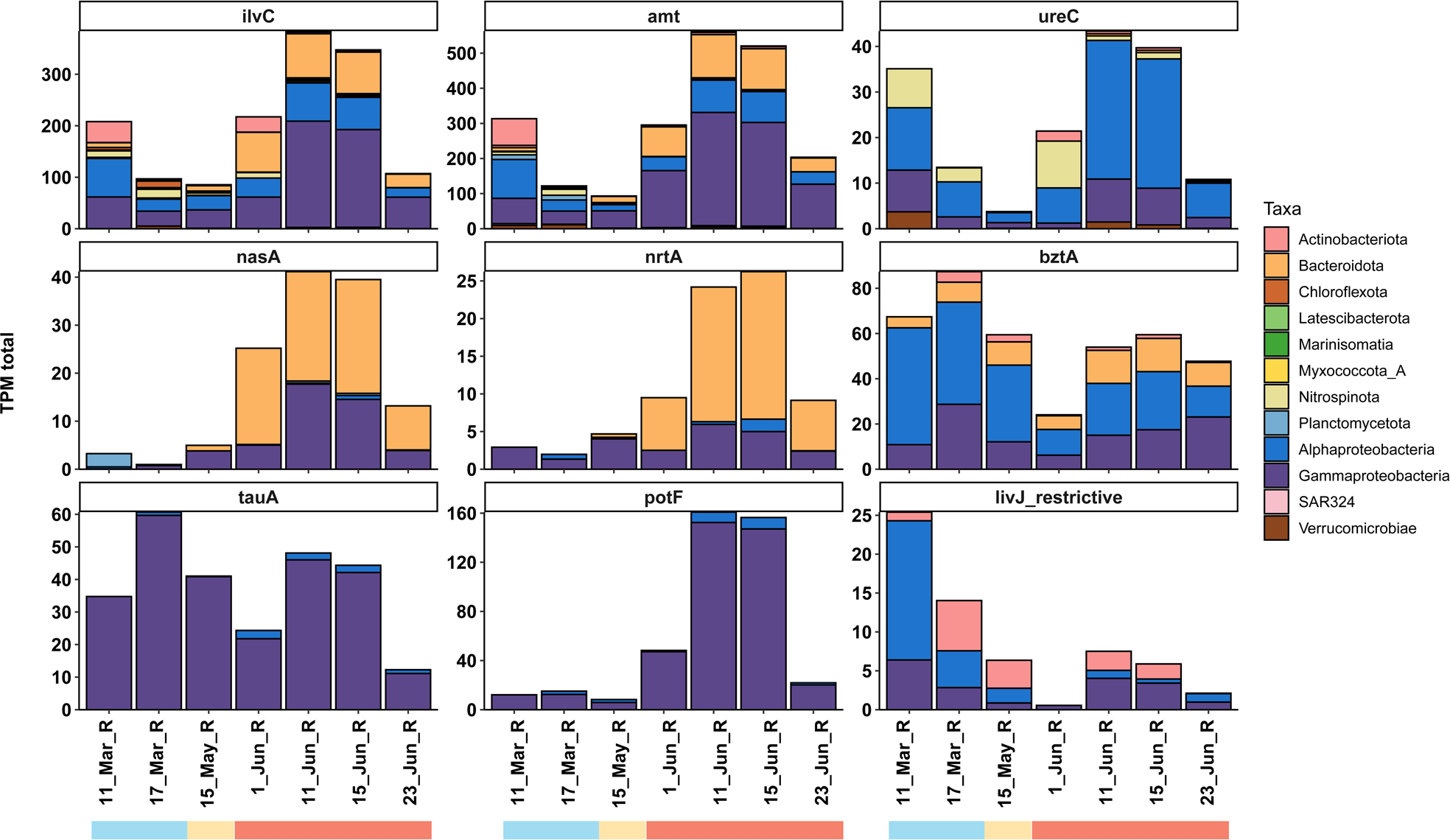
Bacterial expression of different genes related with nitrogen assimilation. Archaeal genes have been removed from the plot for clarity. The colour indicates class taxonomy. Expression of many genes related to nitrogen assimilation seem to increase during summer with the phytoplankton bloom.

**Figure 4.**
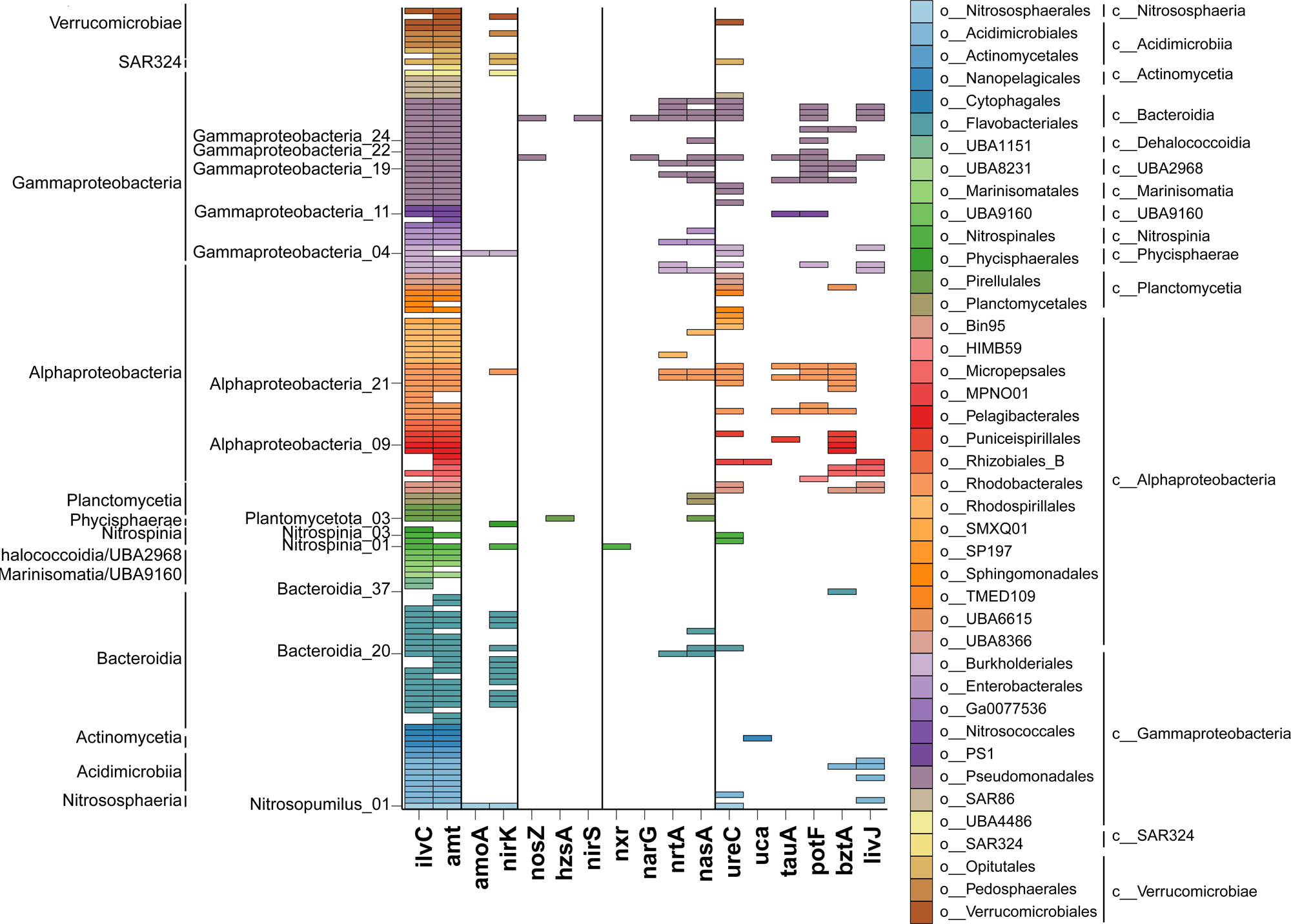
Overview of the nitrogen related genes present in the MAGs. Each row represents a genome and coloured tiles indicate presence of the corresponding gene in the MAG. Colour indicates the taxonomical order. Gene columns are grouped: 1) *ilvC* and *amt* gene that are widespread; 2) *amoA* and *nirK* related to ammonia oxidation; 3) scarce genes; 4) nitrate-cycling genes (*nxr, narG, nrtA, nasA*); 5) genes related to organic nitrogen sources.

#### Nitrate

Nitrate can be the predominant nitrogen source in some Arctic waters according to the *f*-ratio^23^. Nitrate assimilation starts with the transport of nitrate into the cell for which two main transport systems are known: the ABC-type transporters of the NrtABCD/NasDEF group and the MFS-type permease NasA^24^. In this study, we only characterized the ABC-type transporter. Genes encoding the Nrt system were present in 13 MAGs affiliated to the *Alphaproteobacteria, Gammaproteobacteria* and *Bacteroidia* (Figure 4), however expression of the corresponding *nrtA* gene is almost circumscribed to a single MAG during summer (Figure 3), specifically a *Bacteroidia* organism (Bacteroidia_20), which according to genome coverage thrives in June reaching 4-5% relative abundance (Table S2). In fact, this *Bacteroidia* MAG also highly expresses in June a *nasA* gene (Figure 3), which encodes the catalytic subunit of the assimilatory nitrate reductase, indicating that during the summer this organism is actively incorporating nitrate. This nitrate represents a supplementary nitrogen source for this *Bacteroidia* organisms, since an expressed *amt* gene is also present in the genome illustrating how different nitrogen sources are needed to thrive and compete in the context of the phytoplankton bloom. 19 additional genomes from the *Bacteroidia, Gammaproteobacteria, Alphaproteobacteria* and *Plantomycetota* encode for an assimilatory nitrate reductase (nas), in most cases co-occurring with *nrtA* genes in the genome, pointing to a canonical assimilatory function (Figure 4). Again, these additional *nasA* genes show a differential transcription pattern with high expression in June and limited to two organisms: a gammaproteobacterium (Gammaproteobacteria_24) and a *Bacteroidia* genome with three genes encoding NasA proteins (Bacteroidia_21).

#### Taurine

Taurine is an amino acid-like compound of the organosulfonates^25^ group that can serve as a source of energy, carbon, nitrogen and sulfur for marine microorganisms^26,27^. The few studies on the microbial metabolism of taurine in the ocean have shown that taurine metabolism seems to be especially present in heterotrophic bacteria from the SAR11 group^28,29^. In our study, the *tauA* gene, which encodes a subunit of the taurine ABC transporter, was identified in seven MAGs, all *Gammaproteobacteira* or *Alphaproteobacteria* (Figure 4). However, only one of these *tauA* genes from a gammaproteobacterium belonging to the *Thioglobaceae* family (Gammaproteobacteria_11) was transcribed (Figure 3). The expression of this *tauA* gene occurs during the whole studied period suggesting that taurine might be incorporated even outside of the bloom period. The same MAG has genes encoding the ammonia (*amt*) and the putrescine transporters (*potF*), which are only transcribed during the summer (Figures 4 and 5). The potential ability to incorporate three different nitrogen sources (taurine, ammonia and putrescine) might be key for the success of this gammaproteobacterium during the bloom, when it reaches high relative transcriptomic abundances (5-15%). Each strategy might be used depending on the environmental conditions: while taurine incorporation seems to happen during the whole period, transport of ammonia and putrescine only occurs during the bloom, maybe due to increased competition during the bloom or the release of these compounds from dead phytoplankton.

**Figure 5.**
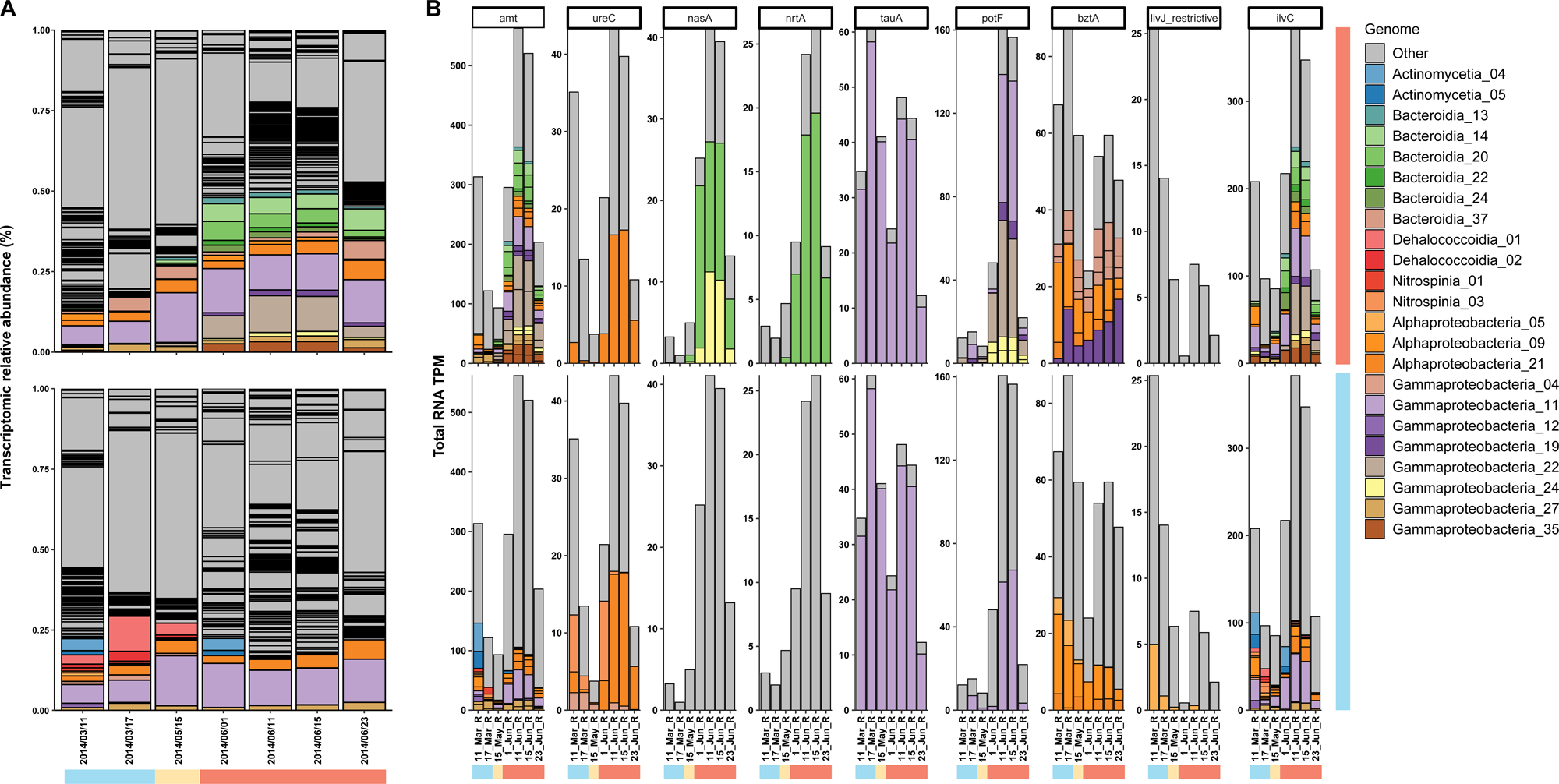
Comparison of the most abundant MAGs of the winter and summer communities. **A** Transcriptomic relative abundances of the different MAGs. Coloured bars indicate the most abundant organisms (with nitrogen-cycling genes) in the sample (top = July; bottom = March), while grey bars indicate other MAGs (or abundant organisms without nitrogen-cycling genes). **B** Transcriptomic profiles for genes related to nitrogen assimilation. Coloured bars indicate the expression of the corresponding gene in the most abundant organisms in the whole dataset (top = July; bottom = March) with nitrogen-cycling genes, while grey bars group the expression of the rest of the genes present in less abundant organisms (or those without nitrogen-cycling genes). The expression profile for some of the genes, i.e. *amt*, is dominated by different organisms indicating the change in the community from winter to summer. There are also higher transcriptomic levels during the summer for many of these genes suggesting a more active nitrogen assimilation.

#### Putrescine

Putrescine is one of the simplest polyamines, organic compounds that consist of multiple amine groups that are naturally produced by many organisms and can serve as carbon and nitrogen source for microorganism^30^. Putrescine has been considered as a model for the cycling of polyamines, which can support up to 10% of the nitrogen needs of the prokaryotic community due to its rapid turnover^31^. In our dataset, 19 MAGs encode for the putrescine ABC transporter (*potF*), which is responsible the translocation of putrescine into the cell (Figure 4). These MAGs are affiliated to the classes *Alphaproteobacteria* and *Gammaproteobacteria*, especially to the order *Pseudomonadales*. However, only two organisms, a *Pseudomonadales* (Gammaproteobacteria_22) and the *Thioglobales* previously mentioned (Gammaproteobacteria_11) showed high expression of the *potF* gene and, moreover, limited to June. This transcriptomic profile links the use of putrescine with the phytoplankton proliferation, likely because phytoplankton is the direct source of putrescine and responsible for its ephemeral increase as previously suggested in other studies^32,33^. Again, it is interesting to note that both MAGs reach high transcriptomic relative abundances (>5%) during the bloom period and possess highly transcribed genes encoding transporters for alternative nitrogen sources like ammonia and taurine, suggesting a fierce competition for nitrogen during the phytoplankton proliferation.

#### Amino acids

To understand utilization of amino acids as nitrogen sources, we studied the presence and expression of two genes encoding the substrate-specific binding proteins of two amino acid transporter systems: the branched amino acid transporter system (LivJ) and the glutamate amino acid transporter system (BztA). Both are ABC transporters, Liv specific for leucine, phenylalanine, isoleucine, valine, and to a lower extent to alanine, threonine and serine, and Bzt for glutamate, glutamine, aspartate and asparagine^34^. In our dataset, different MAGs of *Alphaproteobacteria* and *Gammaproteobacteria*, *Actinobacteriota* and *Bacteroidota* (in this case only for BztA) encode for these systems (Figure 4). Regarding the expression, *bztA* and *livJ* have intermediate expression levels with a similar pattern (Figure 3): highest expression during March, lower transcription in the spring-summer transition and a small increase during June (but not as high as in March). In comparison to other genes, the transcriptomic profiles of *livJ* and *bztA* are more diverse and are not dominated by one or two organisms, indicating that the incorporation of amino acids seems to be a widespread strategy among the prokaryotic community, which is actively used. In general, the *livJ* genes had much lower expression levels than the *bztA*. Still, in both cases most of the organisms expressing *bztA* or *livJ* genes were *Alphaproteobacteria,* like a *Pelagibacterales* MAG (Alphaproteobacteria_09) that dominated the transcriptomic profile of the *bztA* gene with higher expression levels in March, when the MAG transcriptomic relative abundance peaked at 2%. Other organisms with high transcriptomic relative abundances in June (around 1-2%) also had elevated expression of the *bztA* gene at that time like the gammaproteobacterium (Gammaproteobacteria_19) or the flavobacterium (Bacteroidia_37), which actually has three *bztA* genes. *Urea.* Urea is an organic compound with two amine groups that can serve as energy and nitrogen source for microorganisms^35^. There are two microbial mechanisms for the degradation of urea into ammonia. The most widespread involves the three-subunits enzyme urease (UreABC)^36^, while the second route involves two enzymes, urea carboxylase (Uca) being the key component^37^. We could find genes encoding both enzymes in our dataset. The *ureC* gene was much more common (34 MAGs) than the *uca* (2 MAGs). Actually, *uca* genes had almost no expression (Figure 2b), suggesting a minor role in our system. On the contrary, *ureC* genes showed high levels of expressions at two points: in March and in June, in a similar fashion to the expression profiles of other studied genes (i.e. *amt, tauA, bztA*). As mentioned above, the Nitrospinia_03 MAG had the *ureC* gene with the highest expression in March and early June, and might be using urea as a nitrogen and energy source since no gene encoding for an NXR enzyme was found in the genome (Figure 5). In mid-June, one alphaproteobacteria genome (Alphaproteobacteria_21) dominated the *ureC* metatranscriptomic profile, although it also showed considerable levels of *ureC* expression in March. This suggest that this *Alphaproteobacteria* organism might be using urea during the whole period, although with a more intense use during the summer bloom. This may be linked to increased urea concentrations in Arctic waters during the summer^38–40^, probably due to the release of the urea accumulated in the ice during the winter period^41^.

Interestingly, genes encoding a Ure enzyme were present in the Nitrosopumilus_01 MAG. Previous studies have shown that ureases are rare in archaea, but widespread in polar^13^, suggesting that urea might be fuelling nitrification in those environments when ammonia is low or intermittent. Actually, it was shown that urea can serve as substrate for nitrification by *Nitrosopumilus maritimus*^42^. In our study, the *ureC* genes of *Nitrosopumilus* showed almost no expression across all transcriptomic samples, indicating that our *Nitrosopumilus* was likely not using urea as substrate for nitrification.

### Anaerobic nitrogen-cycling processes do not seem to play a role during the bloom

A few genes related to the nitrogen cycle were present in our dataset, but showed almost no expression, suggesting that the associated nitrogen-cycling processes were not relevant during the studied period (Figure 2b). For instance, a *Pseudomonadales* MAG (Gammaproteobacteria_28) harbours genes (*narG, nirS, nosZ*) for a complete denitrification pathway, an anaerobic route where nitrate is used as terminal electron acceptor and is successively reduced to N_2_. This MAG also encodes for different transporters of nitrogen compounds (Amt, NrtA, LivJ, PotF) or the urease enzyme (UreC). However, none of these genes showed high levels of expression. An additional *Pseudomonadales* MAG (Gammaproteobacteria_21) of our dataset had a different *nosZ* gene, but this gene showed no transcription levels at all. Similarly, only one gene of a *Planctomycetia* MAG (Planctomycetota_03) encoded the only hydrazine synthase (HzsA) of this dataset, a marker for the anaerobic oxidation of ammonia. This gene had almost no expression pointing that this process has little relevance in this environment (Figure 2b). The low abundance and expression of the *narG, nirS, nosZ* and *hzsA* genes is not surprising considering that the corresponding enzymes usually function under anaerobic conditions.

## Discussion

We have reconstructed 176 MAGs of an Arctic microbial community from the Canadian Archipelago and studied the expression over four months of different genes encoding nitrogen-cycling enzymes. We observed a change in the community composition from a winter community (March), where chemolithotrophy played a relevant role, to a summer scenario dominated by bloom dynamics with heterotrophic bacteria feeding on phytoplankton-derived carbohydrates and competing for the different nitrogen sources. These metabolic changes were reflected in specific groups and expression profiles. In winter, chemolithotrophic microorganisms were responsible for complete nitrification from ammonia to nitrate: ammonia-oxidizing archaea (*Nitrososphaeria*) converted ammonia to nitrite and then this nitrite was oxidized by *Nitrospinota* bacteria to nitrate. Owing to this metabolic coupling, the corresponding genes encoding the enzymatic machineries showed a similar transcriptomic pattern with a peak of expression in mid-March, declining in May and disappearing by June. This metabolic link explains the concurrence of *Nitrososphaeria* and *Nitrospinota* that has been previously observed in different marine systems^12,19,43–45^.

Previous studies have shown that nitrification rates are highest during winter in Arctic waters^46^. Besides, they have shown that although ammonia has very low concentrations during the whole year, the values tend to be higher during winter^2,46^ when accumulation of nutrients occurs due to the lack of light that prevents primary production, favouring chemolithotrophic lifestyles. Besides, temperature can also influence the rate of nitrification and ammonia assimilation in Arctic microbial communities^46^, explaining why there is higher expression of the *amtA* gene during summer.

With the environmental changes of the Arctic spring/summer, a phytoplankton bloom develops triggering a change in the microbial community. This change is also reflected in the transcriptomic profiles of many organisms and their corresponding nitrogen-cycling genes, where we observe a turning point around May separating two community expression patterns. At that point, the transcription of different machineries for nitrogen assimilation increased in the bacterial community (Figure 2), for example the genes for ammonia (*amt*) and nitrate incorporation (*nrtA, nasA*), urea oxidation (*ureC*) or transporters for organic nitrogen (*potF, bztA*). In general, the expression profiles of these genes were dominated by the most abundant organisms according to transcriptomic relative abundances, which thrive on the phytoplankton-derived organic matter released during the bloom. Many of these highly abundant organisms possessed several ways to incorporate nitrogen, suggesting that part of their success during the bloom might be attributed to their ability to compete for nitrogen. Previous studies have shown that heterotrophic bacteria are key players in the assimilation of different N sources during phytoplankton blooms^47^ and that bacteria proliferating during blooms have more abundant genes for uptake of organic nitrogen compared to other bacteria^48^. Still for some of the most abundant organisms we could only detect the *amt* gene, responsible for ammonia transport, as the only mechanism to incorporate nitrogen. In fact, peaks in ammonia concentration have been reported in earlier studies on Arctic phytoplankton blooms^23^, where it seems that rapid turnover allows that ammonia plays a predominant role as nitrogen source during the bloom^49^. Still, we could not detect any gene for nitrogen incorporation in some organisms, such as in the most abundant organisms through the whole dataset, a *Pelagibacterales,* (Alphaproteobacteria_10). This could be attributed to limitations of the binning process or that these organisms possess other systems that we did not include in the analysis.

The changes in community composition between March and July are also reflected in the expression of nitrogen assimilation mechanisms (Figure 5). In this way, the most abundant organisms in March dominated the expression of nitrogen assimilating genes (like *amt* and *ureC*), but in July other bacteria replace the March community as the most abundant organisms and the ones dominating the expression of mechanisms for nitrogen assimilation. Actually, since June there is an increased expression of additional mechanism for nitrogen incorporation (like NrtA, NasA, PotF) indicating the strong competition for nitrogen compounds and the availability of new nitrogen sources probably linked to organic matter recycling. In general, the highest expression corresponded to the *amt* gene, indicating a preference for ammonia as nitrogen source. The expression of these *amt* genes dramatically increased in June among the bacteria that proliferate during the bloom. Except ammonia transporters, that are widespread, different mechanisms seem to be present in specific groups and especially differentially expressed suggesting a niche specialization. Thus, *Gammaproteobacteria* seem to monopolize organic nitrogen compounds like taurine and putrescine; *Bacteroidetes* are specialized in assimilating nitrate and *Alphaproteobacteria* have higher expression of machineries related to utilization of urea and amino acids (Figure 3). This suggests that specific bacterial groups have the ability to use different nitrogen sources during the summer. This could be interpreted as an intense competition within the community, but also specialization. Future studies should include metabolic and biogeochemical data to support our findings.

## Material and Methods

### Sampling, nucleic acid extraction and sequencing

The sampling campaign, nucleic acid extraction and sequencing were already described in a previous publication from Puente et al. (2022)^50^. In short, samples were collected in Dease Strait, lower Northwest Passage, Nunavut, Canada (69.03°N, 105.33°W) from March until July 2014 within the 2014 Ice Covered Ecosystem-CAMbridge Bay Process Study (ICE-CAMPS). A list of the different samples can be found in the Table S1. DNA was extracted using a modified version of the phenol/chloroform protocol^51^, while RNA was obtained using Qiageńs RNeasy kit. DNA sequencing was performed for most of the samples at CNAG on an Illumina HiSeq2000 sequencing platform using a TruSeq paired-end cluster kit, v3.

### Assembly, binning and metabolic prediction

For each of the metagenomics library, we used SPAdes v3.15.5^52^ to perform a de-novo individual assembly. Afterwards, we performed binning using a combination of Metabat2 v.2.15^59^ and Maxbin v.2.2.7^60^ with a final processing with DasTool v.1.1.2^61^. We used GTDB-tk v.2.1.1^53^ and checkM v.1.1.3^54^ to evaluate the quality and taxonomy of the obtained bins. We combine all the bins from the different individual assemblies and dereplicated them using the dereplication feature of coverM v.0.6.1 (https://github.com/wwood/CoverM) with default settings except the flags ‘--checkm-tab-table -- dereplication-ani 95 --dereplication-prethreshold-ani 90’. Dereplicated bins were further refined via a targeted-reassembly pipeline as described in Laso-Pérez et al.^64^. In short, metagenomic reads were mapped using bbmap (https://sourceforge.net/projects/bbmap/) to the bins. These reads were used for a new assembly with SPAdes v3.15.5 and filtering contigs below 1,500 bp. This procedure was iterated for several rounds for each bin until no further improvement was obtained. We assessed the bin quality using checkM and checkM2 v.1.0.1^55^ by considering completeness, contamination, N50 value and number of scaffolds. Only bins with 50% completeness and contamination below 10% were selected for further analysis and considered as Metagenomic-Assembled Genomes (MAGs). To calculate the MAG relative abundance in our metagenomics and metatranscriptomics samples, we used coverM v.0.6.1 with all the final MAGs. Results were visualized in R v.4.2.1. We used the SqueezeMeta pipeline v.1.5.1^56^ for automatic metabolic prediction of the MAGs and calculation of coverage and abundance estimations of the predicted genes in the different datasets.

### Identification of nitrogen cycling genes and classification

To study the cycling of nitrogen, we developed a database of phylogenetic trees for different marker genes according to the pipeline described in Rivas-Santisteban et al. (2023)^57^ and available in a GitHub repository (https://github.com/Robaina/MetaTag). In this pipeline, high quality trees are constructed to distinguish between true marker genes and closely related paralogues with different functions. Then, candidate genes from the different query MAGs are retrieved using specific protein models (i.e. COGs, arCOGs and KEGG) and they are classified within the previously constructed high-quality tree allowing its classification as true marker genes or a closely related paralogue. We built reference databases for 18 genes encoding different proteins involved in the nitrogen cycle (Table S4). They represent marker genes for different dissimilatory/assimilatory steps of the nitrogen cycle (*amoA, nifH, nirK, ureC, uca, narG, nxrA, nirS, nasA, nosZ, hzsA*), transporter of nitrogen substances (*amt, nrtA, tauA, potF, livJ, bztA*) or biosynthetic pathways (*ilvC*). After tree construction, we searched for the candidate genes in all the MAGs based on the predictions of SqueezeMeta according to specific protein model annotations (Table S4) to then place those query sequences into the phylogenetic trees and distinguish between the true marker genes and closely related paralogues. We kept the true marker genes and extracted their coverage and abundance estimations calculated by the SqueezeMeta pipeline to visualize the temporal and taxonomic changes in our samples by using the R v.4.2.1 software.

## Acknowledgements

We would like to recognize the in-kind support from the Canadian High Arctic Research Station (CHARS) in the Dease Strain sampling. A. Delaforge assisted greatly in data collection and sample processing. Sampling in Dease Strait was funded by an NSERC Discovery and Northern Research Supplement Grant to C.J.M. V. Balagué and M. Royo carried out the nucleic acid extractions at ICM, CSIC. R.L.-P. was funded by a Juan de la Cierva grant (FJC2019-041362-I) from the Spanish Ministerio de Ciencia e Innovación and by a Ramón y Cajal grant (RyC2021-031775-I) from the Spanish Ministerio de Ciencia e Innovación (MCIN/AEI/10.13039/501100011033) and the European Union («NextGenerationEU»/PRTR). Sequencing and data analysis were funded by grant PID2019-110011RB-C33 from the Agencia Estatal de Investigación of the Spanish Ministerio de Economía y Competitividad to J.T. and C.P.-A funded by MCIN/AEI/10.13039/501100011033 and EU PRODIGIO PROJECT GA # 101007006. This research was also funded by projects TRAITS (PID2019-110011RB-C31) of Agencia Estatal de Investigación, Spanish National Plan for Scientific and Technical Research and Innovation.

## Competing Interests

The authors declare no competing interests.

## Data Availability Statement

Generated sequences were deposited under NCBI BioProject ID PRJNA803814. The resulting MAGs were deposited under NCBI BioProject ID PRJNA1044758. During the reviewing process, MAGs and other supplementary material can be found in the following repository: https://saco.csic.es/index.php/s/6Dy84wDF6wNQ2aA

## Table legends

**Table S1**. List of metagenomic and metatranscriptomic samples used in the present study.

**Table S2.** Overview of the metagenome-assembled genomes of our dataset, their assigned taxonomy according to the GTDB database and their relative abundances in the DNA and RNA samples. Their relative abundances are normalized to the mapped reads. Tiles with colour indicate normalized relative abundances over 10% (red), 1% (green) or 0.5% (yellow). Bin quality indicators by chekm and checkm2 are also indicated. Finally, the presence of nitrogen-cycling genes is also shown.

**Table S3.** Overview of the genomic and transcriptomic abundances (in TPM) for the different nitrogen-cycling genes

**Table S4.** List of the studied genes encoding nitrogen-cycling proteins. The corresponding genes were searched in each MAG using the indicated protein motifs.

---

**Figure S1.**
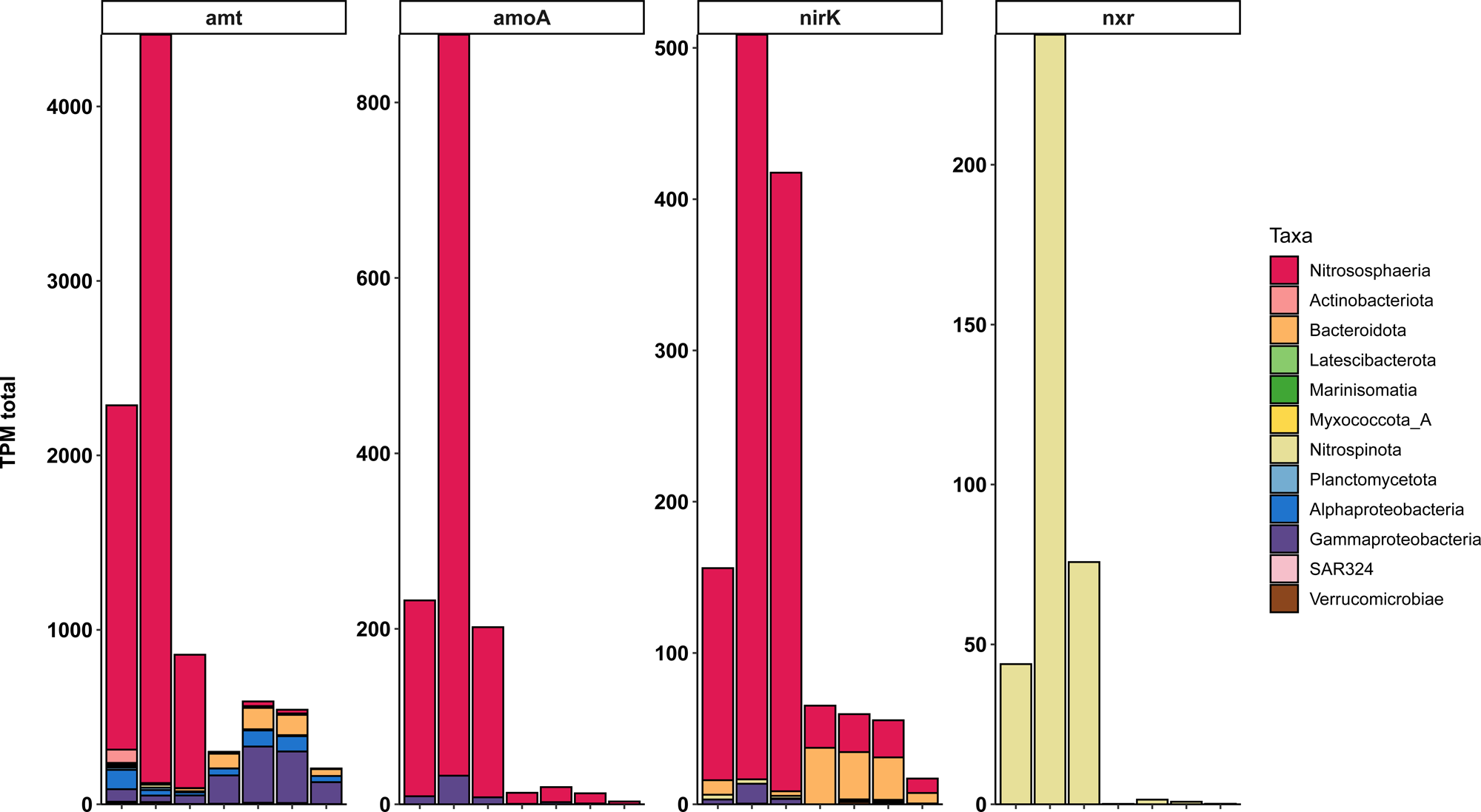
Nitrogen-related chemolithotrophy is present during winter. The four plots represent the RNA expression of the following genes: *amt* (ammonia transporter), *amoA* (ammonia monooxygenase), *nirK* (another step of ammonia oxidation in archaea) and *nxr* (nitrite oxidoreductase). Color indicates class taxonomy. The plots represent the expression for all transcripts of the corresponding genes. In three of them, the genes of the *Nitrososphaeria* MAG dominate the expression profile during winter indicating active ammonia oxidation. As a result, nitrite is produced and the NXR enzyme of the *Nitrospinota* can perform nitrite oxidation. These two processes seem to stop at the summer time.

## Notes

https://saco.csic.es/index.php/s/BpRMcfs6iLQeM9z

### Competing Interest Statement

The authors have declared no competing interest.

